# Host evolutionary history and ecological context modulate the adaptive potential of the microbiome

**DOI:** 10.1101/2020.09.21.306779

**Authors:** Lucas P. Henry, Michael Fernandez, Andrew Webb, Julien F. Ayroles

## Abstract

Can the microbiome serve as a reservoir of adaptive potential for hosts? To address this question, we leveraged ∼150 generations of experimental evolution in *Drosophila melanogaster* in a stressful, high-sugar (HS) diet. We performed a fully reciprocal transplant experiment using the control and evolved HS bacteria. If the microbiome confers benefits to hosts, then transplant recipients should gain fitness benefits compared to controls. Interestingly, we found that benefits do exist, but mismatches between fly evolution and microbiome exerted fitness costs by slowing development and reducing fecundity, especially in the stressful HS diet. The dominant HS bacteria (*Acetobacter pasteurianus*) uniquely encoded several genes to enable uric acid degradation, mediating the toxic effects of uric acid accumulation due to the HS diet for flies. Our study demonstrates that host genotype x microbiome x environment interactions have substantial effects on host phenotype, highlighting how host evolution and ecological context together shape the adaptive potential of the microbiome.

## INTRODUCTION

The microbiome has profound effects on many aspects of organismal biology. From modulating physiology [1] to pathogen protection [2] to social interactions [3], the microbiome is frequently implicated as a key regulator of many host phenotypes--but can the microbiome also influence host adaptation? For hosts where microbes are tightly controlled and faithfully transmitted across generations, microbes can have clear influence on host evolution [4–6]. For example, the acquisition of nutrition-provisioning symbionts enables the ecological expansion of sap-feeding insects [7]. For many eukaryotes, though, their microbiomes are environmentally acquired or through other weakly controlled transmission mechanisms [5]. Weak control over microbial transmission disrupts the links between the microbiome and host over generations. Without these links, the microbiome may have limited effects on host evolution [4, 6, 8]. Microbiomes may still have the potential to influence host evolution because environmentally acquired microbes must survive and adapt to external environments. In doing so, microbiomes may acquire beneficial traits that hosts can leverage. Thus, microbiomes have the potential to precede and facilitate rapid host adaptation, but this process is poorly understood.

Environmentally acquired microbiomes appear to facilitate rapid host adaptation in several systems. For example, bean bugs acquire pesticide resistance from their soil-dwelling *Burkholderia* symbiont [9]. The gut microbiome of woodrats facilitates survival on toxic plant diets [10]. In plants, drought-adapted soil communities can increase fitness in drought conditions [11]. These studies suggest that microbes can adapt and evolve to stressful conditions in natural environments and buffer the stressor for the host, but much about the evolution of these interactions remains unclear—for example, we do not know to what extent these microbial effects depend on host genotypes. Host genetic variation may influence the responsiveness to microbial variation such that some combinations of host and microbiome differentially affect host phenotypes within species. The effects on host phenotype will also likely depend on the environmental context, where mismatches between the environment, host genotype, and microbiome may negatively affect host fitness. In other words, host genotype x microbial genotype x environment (G_H_ x G_M_ x E) interactions may shape how host-microbiome systems evolve. Such complex G_H_ x G_M_ x E interactions are likely to be common, and yet their contribution in shaping host phenotypic variation and adaptation are underexplored. As the study of microbiomes moves beyond descriptive attributes, experimentally assessing the interactions between host and microbial evolution are a key research priority for better understanding the interplay in host-microbiome interactions [12, 13].

The fruit fly, *Drosophila melanogaster*, has emerged as an excellent model to study host-microbe interactions. Flies have relatively simple microbiomes, often composed of fewer than 10 bacterial species from acetic acid and lactic acid bacteria families with effects on many different fly traits [14]. The microbiome is largely environmentally acquired through feeding, defecation, and egg smearing during oviposition [15, 16]. After hatching, larvae acquire their microbes through a combination of microbial persistence and colonization and form associations over the lifespan of the fly [17–19]. Many fly-associated microbes can be individually cultured and reconstituted in various combinations to inoculate sterile flies [14, 20]. With the feasibility of microbiome manipulations and rich genetic resources, *Drosophila* is an ideal model to study the interplay between host genetic, microbiome, and environment (i.e., G_H_ x G_M_ x E interactions) in shaping host phenotypes that underlie rapid host adaptation.

In this study, to investigate whether the microbiome can serve as a reservoir of adaptive potential for hosts, we studied the contribution of the microbiome in the context of experimental evolution. Experimental evolution has a long history of providing fundamental insights into adaptive processes, but has largely neglected the contribution of the microbiome [6, 21, 22]. One particularly powerful approach is the Evolve and Resequence (E&R) framework to measure the genomic response to selection [23]. E&R experiments typically begin with large, outbred populations which are then subjected to a selective regime, and in parallel, control populations are maintained in a benign, non-selective environment. After several generations, the control and evolved populations are then sequenced to identify the genomic response to selection. Although traditionally E&R experiments have focused on the organism targeted by selection (e.g., yeast, flies, mice, etc.), the E&R approach can also identify changes in the microbiome associated with the evolved populations [22]. However, to determine whether any changes in the microbiome over generations of selection actually contributed to host adaptation, it is necessary to transplant the microbiome between control and evolved populations. This approach holds great potential to inform how the microbiome shapes host phenotypes and contributes to host adaptation.

Here, we leverage ∼150 generations of fly and microbial adaptation to a nutritionally stressful, high sugar (HS) diet. HS diets exert stress on flies, and flies exhibit similar obesity-like phenotypes as humans with elevated triglycerides, insulin resistance, and shortened lifespans [24–27]. In turn, the different bacteria in the fly microbiome have a range of effects on fly phenotypes, shaping life history traits [28, 29] and metabolism [28, 30], and these microbiome effects on host phenotype can depend on host genetics [31, 32]. Together, these findings suggest that the microbiome could potentially contribute to host adaptation to the HS diet. Our goal was to determine whether transplanting the adapted microbiome into non-adapted fly genotype would confer adaptive traits in the stressful HS diet. Through a fully reciprocal host x microbiome x diet transplant experiment (Fig. 1), we evaluate how each component contributes to fitness associated traits in flies. The power of our approach is to compare the benefit of adapted microbes relative to non-adapted microbes under different host genetic and environmental contexts. Our results point to the important role microbiomes may play in influencing host adaptation but also highlight the complexities of environmental context dependence on host-microbiome interactions.

**Figure 1:**
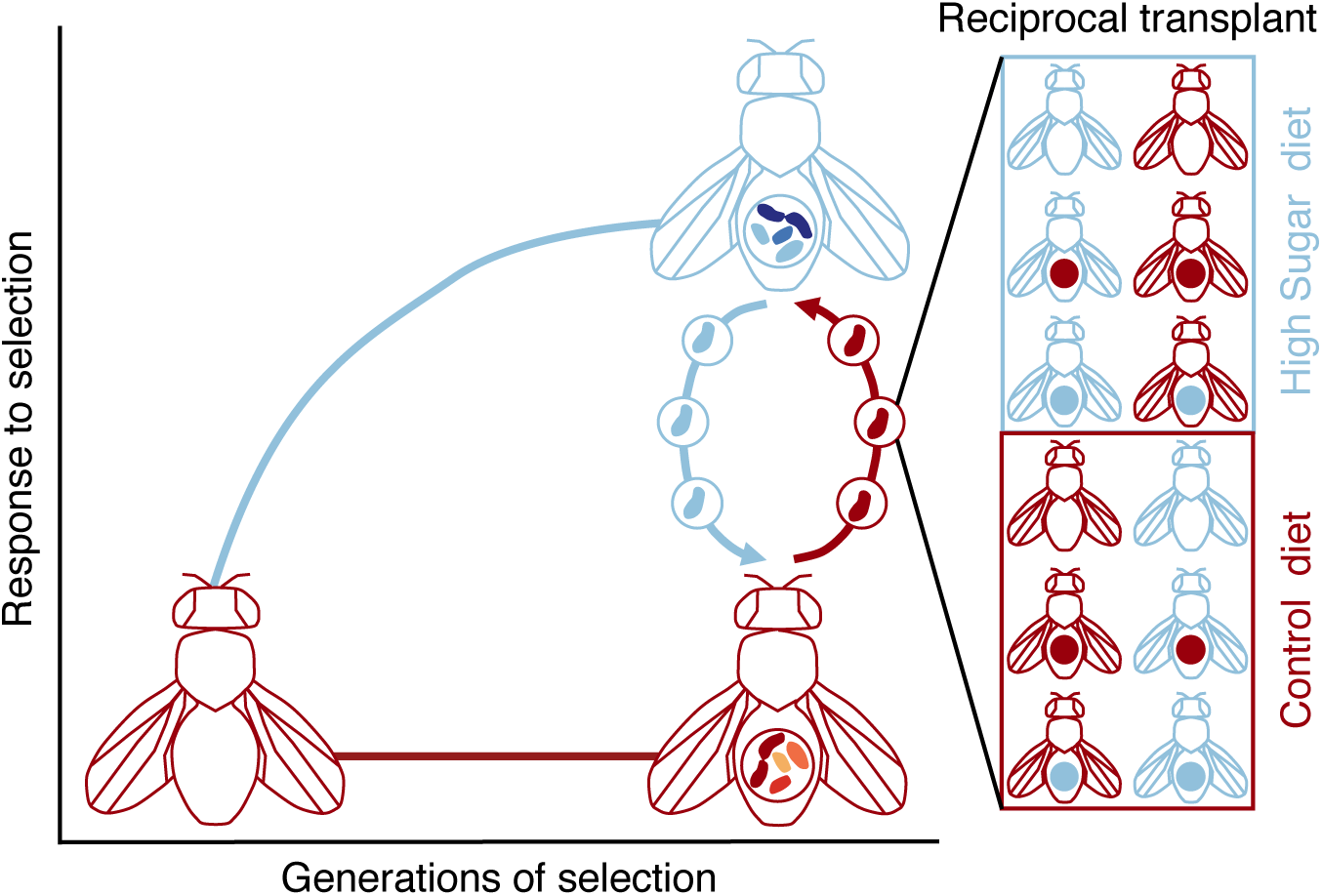
Experimental design to leverage ∼150 generations of fly adaptation to the high sugar (HS) diet to study the adaptive potential of the microbiome. The conceptualized evolutionary trajectory shows control (C) flies and microbiome in red and HS flies and microbiome in blue. By the end of experimental evolution, the C fly population (red) has a different microbiome than the HS fly population (blue). After the ∼150 generations of selection, we performed a fully reciprocal host x microbiome x diet transplant experiment for all combinations of fly genotypes (C or HS), microbes (sterile, C, or HS), and diets (C, or HS). Together, this approach allows for the assessment of the benefit of adapted relative to non-adapted microbes in both host genetic and environmental contexts.

## MATERIALS AND METHODS

### Experimental design

We first identified if the microbiome responded to selection from experimental evolution to the HS diet. Then, to test if the microbiome can transfer adaptive traits, we performed a reciprocal microbiome transplant experiment (Fig. 1). All host genotypes x microbiome x diet interactions were generated in the reciprocal transplant. Both genotypes (C and HS flies) were inoculated with microbiomes (C, HS) or kept sterile and raised in both diets (C and HS) for a total of 12 treatments. This experimental design allowed us to evaluate how host x microbiome x environment interactions influence the transfer of adaptive phenotypes.

### Fly populations

All fly populations were maintained at 25°C with 12 hour light:dark cycles. The base ancestral population was derived by round-robin crossing a subset of the Global Diversity lines [33], which were then maintained at large population size (>10,000 flies) and allowed to interbreed freely for ∼50 generations before HS selection began. The control (C) diet was composed of 8% glucose, 8% yeast, 1.2% agar, with 0.04% phosphoric acid and 0.4% propionic acid as preservatives. Three replicate populations for HS selection were derived from the base population and maintained on the same diet, but with an additional 12% sucrose and at lower population size (>5,000 flies/cage). The control population was the ancestral population maintained at large population size (>10,000 flies) in a parallel on the C diet during the entirety of the HS selection. The approach used here allowed for natural selection on HS diets; we did not guide or select for particular traits, which is a common approach in experimental evolution studies in *Drosophila*. We consider the two populations, C and HS, to be two different genotypes, though note they are outbred and retain genetic diversity.

### Identifying the microbiome response to selection in the HS diet

We first identified how the microbiome changed between C and HS flies through 16S rRNA amplicon sequencing. In brief, DNA was extracted from single, age-matched (7-10 days post eclosion adults) using Zymo Quick-DNA extraction kit (Zymo D3012). The 16S V1-V2 region was amplified, pooled, and digested with BstZ17I (NEB R3954) to deplete Wolbachia reads [34]. Libraries were sequenced using 300 bp paired-end reads using the Illumina MiSeq platform at the Princeton University Genomics Core. We used QIIME2 v2019.4 [35] to process reads, cluster into amplicon sequence variants (ASVs) with DADA2, and assign taxonomy with Greengenes classifier trimmed to the 16S rRNA V1-V2 regions. Data was imported into phyloseq for visualization [36]. Flies with fewer than 500 reads/individual were discarded before the analyses.

To identify differentially abundant bacteria between the C and HS flies, we used ANCOM (analysis of composition of microbiomes) [37] implemented in QIIME2. We cultured the most differentially abundant bacteria from HS flies, *A. pasteurianus*, and the significantly differentially abundant bacteria within the *Acetobacter* genus and enriched in C flies, *A. indonesiensis*. We confirmed 16S rRNA sequence identity using Sanger sequencing. For simplicity, we refer to *A. indonesiensis* as C *Acetobacter* and *A. pasteurianus* as HS *Acetobacter*.

### Transplanting microbiomes to test for adaptive significance

We first performed both a complete microbiome transplant. We then performed a focused transplant of the dominant C and HS *Acetobacter* strains identified through ANCOM to measure the impact of host genetic, microbial, and environment interactions on fitness-associated phenotypes.

First, to transplant the complete microbiome, we collected frass for ∼ 5 hours from 300 C and HS donor flies (age matched 7-10 days post eclosion). Flies were placed in sterile 50 ml tubes, lined with a plastic sheet. Frass was then washed off the plastic using sterile water and filtered through fine mesh to remove particulates and any eggs. Preliminary tests suggested that the HS microbiome had much lower bacterial abundance, and so the C microbiome was resuspended in 6 ml sterile water and 3 ml for the HS microbiome. 50 µl of the frass wash was used to inoculate axenic eggs in sterile diets following standard protocols [20], and flies were reared in a biosafety cabinet at 25°C and 12:12 light:dark cycle through the lifespan of the recipient flies. At 7-10 days post eclosion, recipient flies were stored at −80°C and DNA subsequently extracted from individuals. The microbiome of recipients was assessed through 16S rRNA V1-V2 profiling as described above and rarified to 500 reads/fly. To assess the fidelity of the microbiome transplant, we tested for the effects of host genotype and microbiome on Bray-Curtis dissimilarity for each diet separately using the ADONIS2 implementation of PERMANOVA in vegan [38].

To perform the *Acetobacter* transplant, both *Acetobacter* strains were cultured in liquid MRS, density was normalized to OD_600_ = 0.1, and 50 µl of bacteria (or PBS for sterile treatments) was used to inoculate axenic eggs in sterile diets [20]. For each treatment, 4-12 replicate populations were maintained. Flies were then reared in a biosafety cabinet at 25°C and 12:12 light:dark cycle through the lifespan of the recipient flies. For the *Acetobacter* transplant, we measured two fitness-associated phenotypes: developmental time and fecundity.

To measure developmental time, we counted the number of new pupae every 6 hours during their daytime (i.e., 8am, 2pm, 8pm) until all treatments began eclosing (4-12 replicates/treatment, see Supp. Table 1 for sample sizes and replication). For developmental time, we used the survival [39] and survminer [40] packages in R to analyze development data. We used Cox proportional hazards to identify how interactions between fly genotype, microbiome, and diet influence development time.

For fecundity, we counted the number of eggs laid by individual 8-10 day old females (N=644 total, see Supp. Table 2 for sample sizes). Females were placed in a 24-well plate with oviposition media. Oviposition media was the same as the fly food media, but with yeast extract, which makes the diet transparent and easier to count eggs. Females laid eggs on both C and HS media; media did not affect egg lay (Mann-Whitney-Wilcoxon test, W=50964, p=0.09), and data were pooled from both oviposition media for analysis. To minimize plate effects, two treatments were assayed on each plate, and treatments were randomly assigned across plates. Females laid eggs for ∼4-6 hrs at 25°C during the late afternoon (2pm-8pm) to capture similar circadian rhythms in egg laying across all treatments. We used a zero-inflated Poisson distribution implemented in glmmTMB [41] testing for the effects of interactions between fly genotype, *Acetobacter* strain, and diet with oviposition plate (N=29 plates) as the random effect.

### Characterizing the genomic differences between C and HS Acetobacter

To further characterize the mechanisms underlying the fitness-associated benefits from the C and HS *Acetobacter* strains, we performed whole genome sequencing on these bacteria. Libraries were prepared following the manufacturer instructions using the Illumina PCR-free Library Prep kit, and then 150-bp paired end reads were sequenced on an Illumina NovaSeq platform. Genomes were assembled using SPAdes [42] with --isolate flag and annotated using the Rapid Annotation using Subsystem Technology (RAST) server [43].

We first compared average nucleotide identity (ANI) between the C and HS *Acetobacter* using OrthoANIu [44]. OrthoANIu computationally fragments bacterial genomes into 1020 bp (contigs <1020 bp are discarded), performs reciprocal BLASTn searches for all fragments, and then computes ANI for all reciprocal BLASTn hits. Then to identify functional changes, we manually compared the RAST annotations to identify pathways unique to either C or HS *Acetobacter*. We also generated the RAST annotations from two other abundant bacteria in the C microbiome and compared them with the HS *Acetobacter* annotations: *Algoriella xinjiangensis* (NCBI accession FOUZ01000040.1) and *Acetobacter persici* (NCBI accession JOPC00000000.1). To confirm the differences between C and HS *Acetobacter* strains, we identified orthologs between the two *Acetobacter* genomes using OrthoFinder v2.5.2 [45]. Then to gain insight into the function of the putative genes, we performed BLASTP v2.10.1+ searches against the NCBI non-redundant protein sequence database [46].

The genomic analysis performed above suggested that the HS *Acetobacter* genome encoded several pathways for urea degradation, but were not present in the C *Acetobacter* genome. To validate this observation, we measured the ability of the *Acetobacter* strains to degrade uric acid in fly food. Uric acid was added at 50 µmol concentration to 30 µl of C and HS diet. Then 10 µl of C or HS Acetobacter (or sterile PBS control), normalized to OD_600_=0.1, were added to the diet and incubated for 4 days at 25°C to degrade the uric acid. Uric acid concentration was measured using the Amplex Red Uric Acid/Uricase kit (ThermoFisher A22181) following the manufacturer instructions. ANOVA was used to analyze the uric acid degradation by bacteria. We tested for the effects of bacteria (C or HS *Acetobacter* strain), diet, and the interaction between bacteria and diet.

## RESULTS

### Fly microbiome responds to HS selection

First, we examined if the microbiome differed between C and HS flies following ∼150 generations of adaptation to the C or HS diets (Fig. 2A). HS flies were dominated by *Acetobacter pasteurianus* (86.7% +/- 1.8 SE) and unclassified Acetobacteraceae strain (5.1% +/- 1.8 SE), while C flies were more diverse with *Algoriella xinjiangensis* (37.3% +/- 5.3 SE), *Acetobacter persici* (16.9% +/-2.6 SE), and *Acetobacter indonesiensis* (14.8% +/- 2.1 SE). The most differentially abundant bacteria were *A. pasteurianus* and Acetobacteraceae strain for the HS flies, while *A. xinjiangensis* and *A. indonesiensis* were associated with C flies (Fig. 2B, Supp. Table 3). These bacteria are unique to either C or HS flies and found across individual flies, demonstrating that the HS diet selected for a different fly microbiome, dominated by *A. pasteurianaus*.

**Figure 2:**
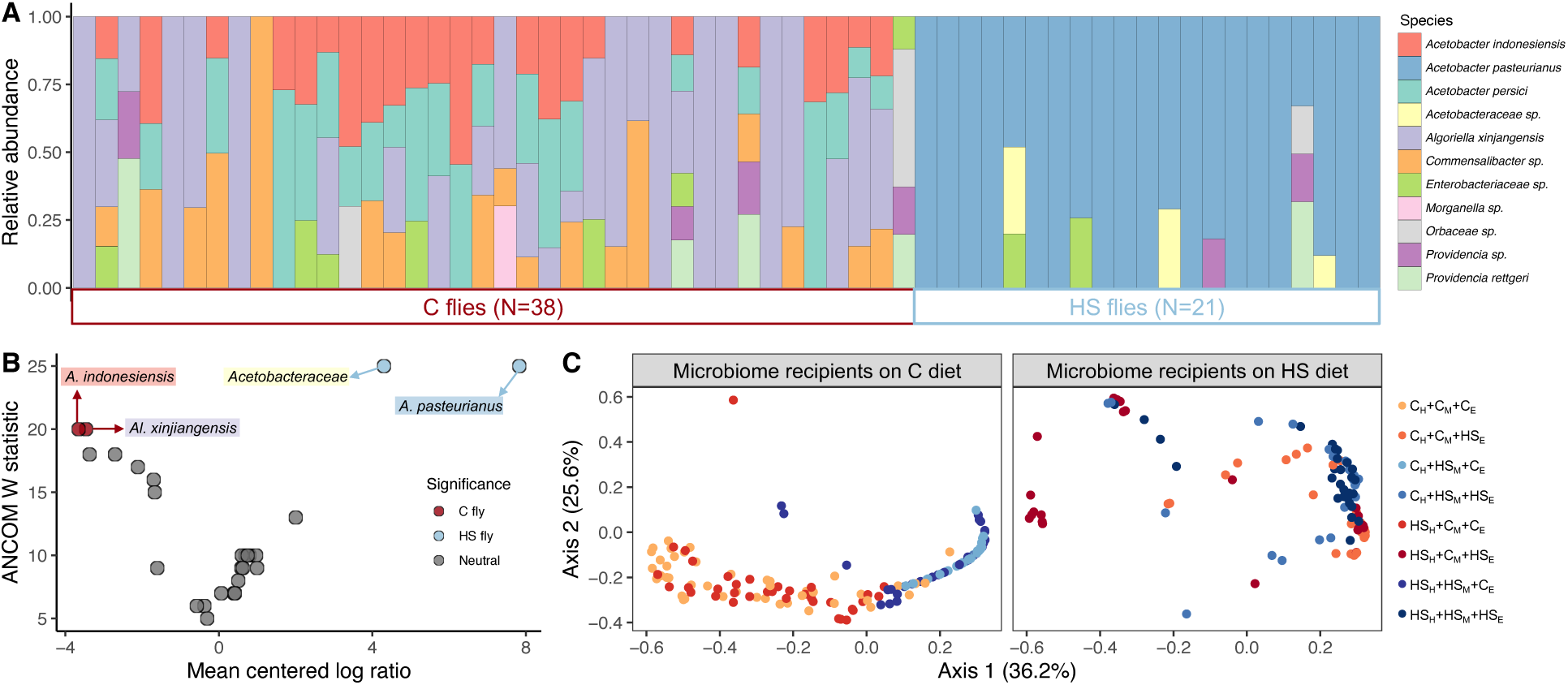
The microbiome in HS flies is different from C flies. A) Relative abundance of bacteria determined using 16S rRNA sequences. Each bar represents an individual fly (N=38 C flies, 21 HS flies). The C fly microbiome is enriched for *A. indonesiensis*, *A. persici*, and *Algoriella xinjiangensis*. The HS fly microbiome is enriched for *A. pasteurianus* and unclassified *Acetobacteraceae*. B) ANCOM results for differentially abundant bacteria between C and HS flies. Each point represents a different ASV, colored by whether significantly enriched for C (red) or HS flies (blue). Labels denote the taxonomic classification for significant ASVs and are highlighted to match the color from the relative abundance in 2A. C) PCoA plots based on Bray-Curtis dissimilarity for recipients in the whole microbiome transplant. Plot is faceted by diet, and colors represent the different combination of host genotype, microbiome, and diet. The difference between C and HS microbiomes was maintained for recipients on the C diet, but not on the HS diet.

### Transplanting Acetobacter influences fitness-related traits

To test if transferring the microbiome could transfer adaptive phenotypes to naive hosts, we performed a fully reciprocal, host genotype x microbiome x diet transplant experiment. We first transplanted a complete microbiome. On the C diet, the difference in C and HS microbiomes were maintained in recipients (Fig. 2C, Supp. Table 4, PERMANOVA, R^2^ = 0.38, p = 0.001). However, the microbiome was not successfully transplanted for flies on the HS diet, as the variance explained by microbiome was much lower in recipients (Fig. 2C, Supp. Table 5, PERMANOVA, R^2^ = 0.04, p = 0.004). This suggests that the HS diet was highly selective and as we discuss later, likely limits bacterial survival.

Instead, given the simplicity of the microbiome associated with HS flies, we performed another microbial transplant using the *Acetobacter* strains identified as differentially abundant between C and HS flies: *A. indonesiensis* (C *Acetobacter*) and *A. pasteurianus* (HS *Acetobacter*). While not the complete bacterial community from the C flies, we reasoned that because both community complexity (i.e., single dominant species in the HS microbiome versus several in the C microbiome) and phylogenetic diversity (i.e., single dominant *Acetobacter* in the HS microbiome versus two *Acetobacter* species, *Algoriella*, *Commensalibacter*, etc. in the C microbiome) differ, using only C *Acetobacter* still provides an ecologically relevant substitute, while controlling for differences in diversity between the C and HS microbiome. We then measured higher order, fitness-associated traits (developmental timing and fecundity), which reflect cumulative adaptations to the multitude of selective pressures in the HS diet.

Development was shaped by interactions between fly genotype, *Acetobacter*, and diet (Fig. 3A, Supp. Table 6). On the C diet, fly genotype influenced the response to the *Acetobacter* transplant. HS flies with the C *Acetobacter* developed more rapidly (avg. 140.8 hrs +/- 0.54 SE) than with the HS *Acetobacter* (avg. 147.6 hrs +/- 0.60 SE). There was no effect for the C fly. However, on the HS diet, the HS *Acetobacter* accelerated development (HS *Acetobacter* avg. 176.5 hrs +/- .50 SE, C *Acetobacter* avg. 186.8 hrs +/-0.60 SE). The three-way interaction between fly genotype, *Acetobacter*, and diet significantly affected development time (log-likelihood =-58500, X^2^ = 47.625, p < 0.0001).

**Figure 3:**
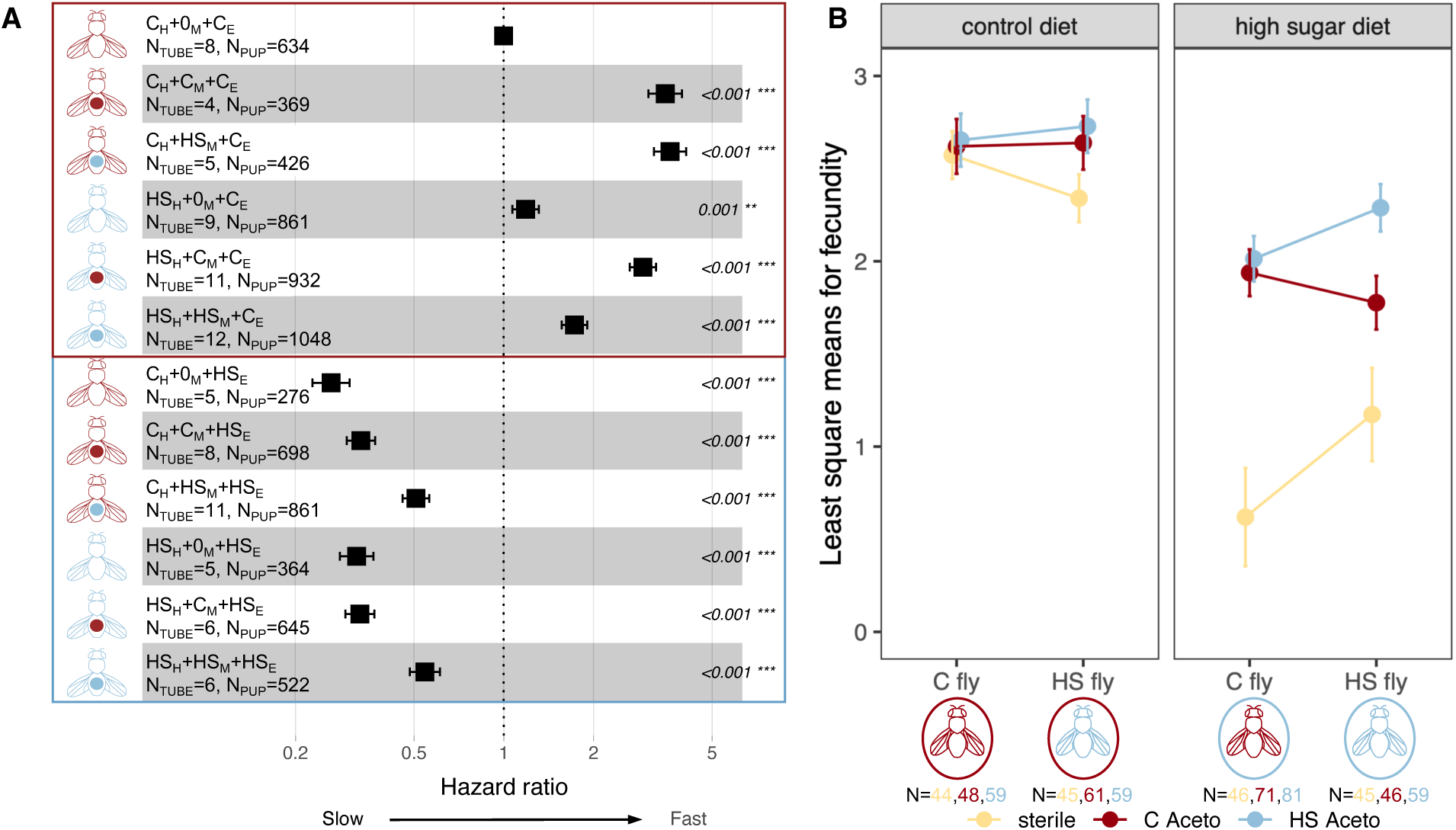
Effects of the *Acetobacter* transplant on fitness-related traits. A) Hazard ratio plot for *Acetobacter* effects on development time. Treatment is shown with the fly diagram, where fly color represents fly genotype, circle represents *Acetobacter* (no circle = sterile), and box represents diet. The full treatment is written out (i.e., C_H_+C_M_+C_E_ = C fly with C microbiome on the C diet) with the total number of replicates (N_TUBE_) and pupae (N_PUP_) phenotyped across replicates shown below. Boxplots show hazard ratios and 95% confidence intervals for each treatment relative to C_H_+0_M_+C_E_. Mismatches between diet and microbiome slowed development. In the C diet, HS flies with C *Acetobacter* developed faster than the HS flies with HS *Acetobacter*. However, on the HS diet, C *Acetobacter* slowed, while HS *Acetobacter* accelerated development. B) Least square means (LSM) from fecundity, modeled with a zero-inflated Poisson distribution show fly genotype x *Acetobacter* x diet interactions shaped fecundity. Plots are faceted by fly diet, where colors represent *Acetobacter* treatment, and the number of age-matched females assayed below the fly diagram. Error bars show standard error for LSMs. For the HS fly genotype on the HS diet, the HS *Acetobacter* increased fecundity relative to the C *Acetobacter*, suggesting a fecundity benefit with the adapted microbiome.

Fecundity responded to interactions between fly genotype, *Acetobacter*, and diet (Fig. 3B, Supp. Table 7). On average, treatments were more fecund on the C diet, while treatments on HS diets laid 54.3% fewer eggs (C avg. 13.0 eggs +/-0.6 SE, HS, avg. 5.9 eggs +/- 0.4 SE). Sterile treatments substantially lowered fecundity, only on the HS diet (+microbes avg. 7.45 eggs +/- 0.5 SE, sterile avg. 1.67 eggs +/- 0.4 SE). Notably, the response to *Acetobacter* treatment depended on fly genotype for the HS diet. On the HS diet, HS flies with HS *Acetobacter* (avg. 8.4 eggs +/-1.4 SE) were almost twice as fecund as flies with the C *Acetobacter* (avg. 4.5 eggs +/- 1.0 SE). The interaction between fly genotype, microbiome, and diet significantly affected fecundity (interaction X^2^ = 16.4, df = 2, p=0.0003).

### HS Acetobacter uniquely encodes uric acid degradation pathway

To better understand how the HS *Acetobacter* may benefit flies in the HS diet, we sequenced and compared the genomes of the C and HS *Acetobacter* strains. ANI between the strains was 73.68% (Supp. Table 8). Annotation by RAST showed that strains shared a majority of predicted functions (>90% for both strains, Supp. Fig. 1). One key difference between the two genomes was that HS *Acetobacter* encodes for several genes that degrade uric acid, but C *Acetobacter* does not (Supp. Table 9). Further, the other dominant bacteria in the C microbiome (*Algoriella xinjiangensis* and *Acetobacter persici*) also lack uric acid degradation genes (Supp. Table 9). Indeed, we found that interactions between *Acetobacter* and the diet shape uric acid degradation (Fig. 4, Supp. Table 10). Only on the HS diet does HS *Acetobacter* reduce the concentration of uric acid by ∼50% compared to C *Acetobacter*, with only negligible changes to uric acid concentration on the C diet.

**Figure 4:**
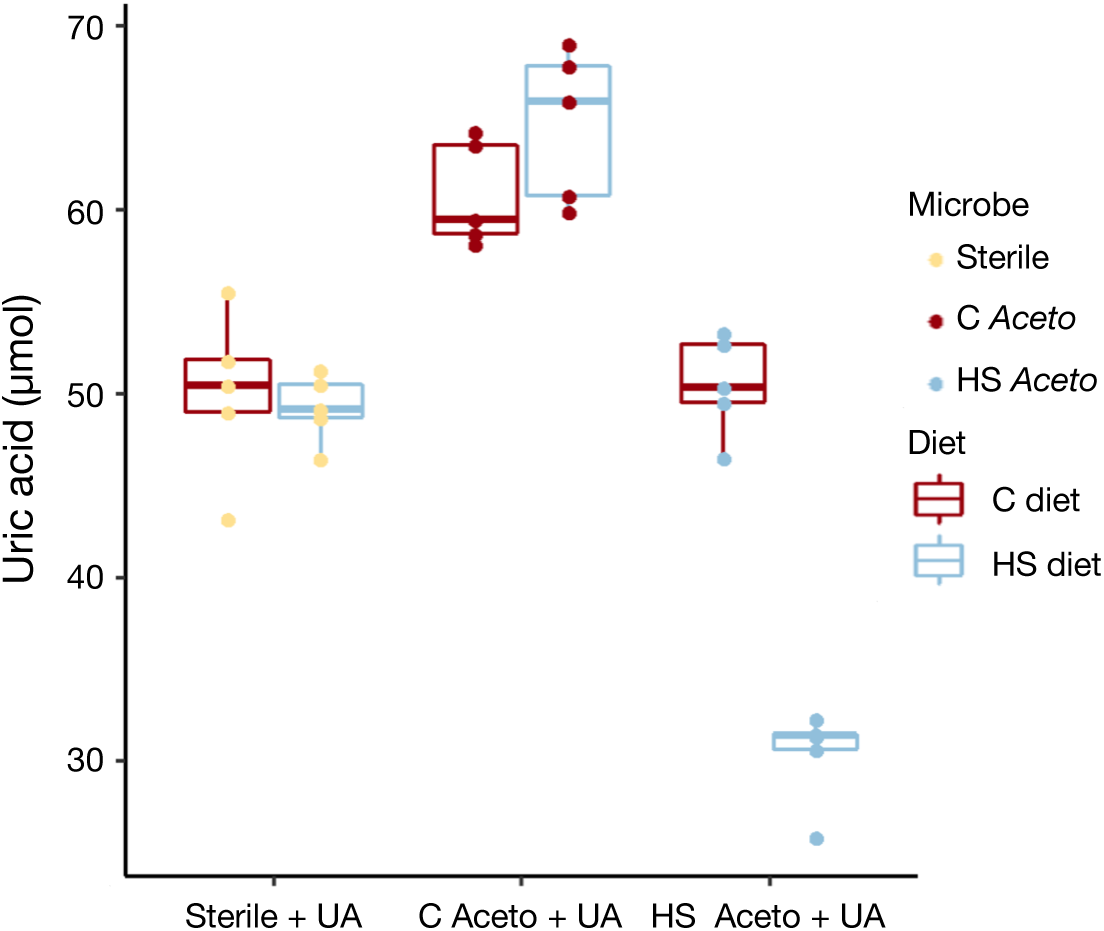
Uric acid degradation by HS *Acetobacter*. Boxplots show uric acid in the *in vitro* diet assay. Boxplots are colored by diet with points colored by *Acetobacter* treatment. Only in the HS diet, HS *Acetobacter* degraded significantly more uric acid than C *Acetobacter*.

## DISCUSSION

Here, we explored how the microbiome influences host fitness in stressful environments. Because all combinations of G_H_ x G_M_ x E interactions were produced in the transplant experiment, the relative benefits of C and HS *Acetobacter* across the different host genetic (selected or unselected) and environmental conditions (HS or control diet) could be evaluated. Through reciprocal host x microbiome x diet transplant experiments, we show that G_H_ x G_M_ x E interactions shape host fitness by modifying developmental timing and fecundity. The HS *Acetobacter* genome encoded uric acid degradation genes and, *in vitro*, lowered uric acid on the HS diet, while the C *Acetobacter* did not. Our results highlight the complex interactions generated by the microbiome, and we discuss important future directions to further dissect these relationships.

### Acetobacter dominates HS microbiome

The HS microbiome was dominated by a single species of bacteria, *Acetobacter pasteurianus* (Fig. 2). *Acetobacter* species are common in flies, both in laboratory settings and wild environments [18, 47–51]. While *Acetobacter* are typically sugar specialists [52], one major axis of variation separating the C and HS microbiome was the ability to degrade uric acid. We did not find an effect of C or HS *Acetobacter* strains in shaping triglyceride accumulation (Supp. Fig. 2); triglyceride accumulation is often a benefit associated with *Acetobacter* in *Drosophila* [28, 31]. In the context of *Drosophila* biology, uric acid accumulation is commonly associated with high sugar diets in flies [53]. The accumulation of uric acid may also exacerbate the deleterious effects of HS diet, altering oxidative stress and inflammation that can promote obesogenic phenotypes [54, 55]. Uric acid is normally excreted by flies, but high concentrations slow development [56] and shorten lifespan [53, 57]. Not all *Acetobacter* species can degrade uric acid, including the C *Acetobacter*, but the ability to degrade is more common in fly-environment than in other free-living *Acetobacter* species [55]. Here, comparative analyses between the C and HS *Acetobacter* genomes showed that HS *Acetobacter* encoded for several genes in the uric acid degradation pathway, while the C *Acetobacter* genome did not. Uric acid degradation may be the key fitness benefit of the HS *Acetobacter* over the C *Acetobacter* in the HS diet.

The fitness advantage of uric acid degradation was likely direct for the HS *Acetobacter*, but indirect for the fly. Over the course of experimental evolution, both fly and microbiome were exposed to the HS diet, and in turn, the HS microbiome was dominated by *A. pasteurianus*, which could degrade uric acid. *Acetobacter* species that degrade uric acid can outcompete *Acetobacter* species that cannot in the fly environment [55]. For the HS *Acetobacter*, uric acid degradation may underlie its persistence in the stressful HS diet, and thus indirectly benefiting flies through reductions of uric acid in the diet. The benefit of uric acid degradation may be especially important during development. Larvae have limited endogenous abilities to detoxify uric acid, and the primary mechanism of uric acid tolerance is by decreasing exposure through decreased uptake or increased excretion [56, 58, 59]. For adults, it is unclear if HS *Acetobacter* also degrades uric acid in the fly gut or simply benefiting from the reduction in the diet. While these questions require further characterization of fly physiology, uric acid degradation by HS *Acetobacter* is similar to a handful of examples of environmentally acquired microbiomes that facilitate adaptation through detoxifying the environment, such as insecticide resistance in bean bugs [9] or terpene detoxification in woodrats [10]. Our results contribute to a growing body of literature that suggests that the microbiome can encode novel functions that hosts can leverage the microbiome during adaptation to stressful environments [6].

### Adaptive potential of the microbiome depends on evolutionary and ecological context

Many have predicted that the microbiome could accelerate local adaptation in the host, facilitating phenotypic changes to buffer local stressors [6, 7, 60, 61]. If so, transplanting the locally adaptive microbiomes should confer adaptive phenotypes to naive individuals. However, the benefit may depend on interactions between host genotype and environment. G_H_ x G_M_ x E interactions have the potential to facilitate or impede evolution in challenging environments [12, 13]. Through combining experimental evolution with reciprocal transplants, our data allowed us to test if and how ecological context (i.e., diet) and evolutionary context (i.e., fly genotype) shape the adaptive potential of the microbiome.

To shape adaptive potential, the microbiome must contribute to host phenotypic variation, not unlike traditional (host) additive genetic variance [6]. The G_H_ x G_M_ x E interaction for development illustrates how diet (the ecological context), when matched with host genotype (the evolutionary context) may increase the fitness benefit of locally adaptive microbiomes (Fig. 3A). The HS *Acetobacter* accelerated development on the HS diet (a match between microbe and diet, independent of fly genotype) and slowed development compared to the C *Acetobacter* on the C diet for only the HS fly (a mismatch between microbe and environment, but dependent on fly genotype). The mismatch only mattered for the HS fly; C fly development was similar between the C and HS *Acetobacter* on the C diet. However, the G_H_ x G_M_ x E interaction that shaped fecundity is suggestive of how genetic interactions may impede the transfer of adaptive potential. The naive host, C flies, did not receive a fecundity increase from the HS *Acetobacter* in the HS diet, but HS flies did (Fig. 3C). The presence of G_H_ x G_M_ x E interactions for both development and fecundity suggests that the microbiome contributed to fly evolution. HS flies responded differently to the *Acetobacter* variation from the C flies, and more so, responsiveness depended on the diet. Because host genotypes may vary in their ability to leverage beneficial microbes in stressful environments, genetic interactions may limit the benefits of locally adaptive microbiomes. Understanding these G_H_ x G_M_ interactions will be especially important in using evolution to engineer beneficial microbiomes to solve pressing problems in public health and agriculture [6, 12, 13, 62]. While longitudinal sampling of the evolutionary trajectory is necessary to determine to what extent the HS fly evolved or selected for beneficial *Acetobacter* traits, our results clearly demonstrate how the ecological context and host evolution can shape the adaptive potential of the microbiome.

### Research priorities

Here, we leveraged host and microbial evolution from ∼150 generations of adaptation to a stressful diet through the E&R approach. Such experiments have illuminated many important findings about the role of demography, population genetics, and genetic architecture in shaping adaptive traits [23, 63]. However, missing from this framework has been the role of the microbiome. In a recent survey of ten E&R experiments, the microbiome also frequently responded to experimental evolution, particularly in traits associated with metabolism [22]. Taken together with our reciprocal transplant experiment, our data also highlight how the microbiome may contribute to host adaptation – effects previously unseen when focusing on the *Drosophila* genome. Recent work on the *Drosophila* microbiome linked to natural variation in wild populations suggested that a single inoculation of two different bacteria linked to different life-history strategies, can lead to rapid phenotypic change and genetic differentiation in one fly population [29, 64]. Our current study advances these findings by comparing differences in ecologically relevant *Acetobacter* evolution to differences in fly evolution, providing one of the first experimental dissections of host-microbiome interactions in the E&R context. Through reciprocal transplants, we identified how interactions between fly genotype, microbiome, and diet shape fitness related traits.

The G_H_ x G_M_ x E interactions observed here highlight how evolution in both the host and microbiome occurred during adaptation to the HS diet. In particular, the ability of HS *Acetobacter* to degrade uric acid in the HS diet reflects the potential for diffuse coevolution [65]. The HS diet likely led to an increase in uric acid production by flies, and then the *Acetobacter* that could exploit the new niche increased, which the HS flies benefited from the reduction in toxicity. Ultimately, in our system and perhaps many others, the evolutionary benefits between host and microbiome are intertwined. Future work utilizing experimental evolution will demonstrate how the joint evolution of host and microbiome shapes the response to selection, providing fundamental insights into how natural selection operates on complex systems.

## ACKNOWLEDGEMENTS

We thank members of the Ayroles lab for helpful feedback on this work. LPH was supported by NSF-GRFP under grant DGE1656466 and National Institutes of Health (NIH) grant R35-GM124881 to JFA.

## DATA AVAILABILITY

Genome sequencing will be uploaded to NCBI [upon acceptance]. Phenotypic data will be deposited in Dryad [upon acceptance]. Code used to analyze data will be posted on github [upon acceptance].

---

**Supp. Fig. 1:**
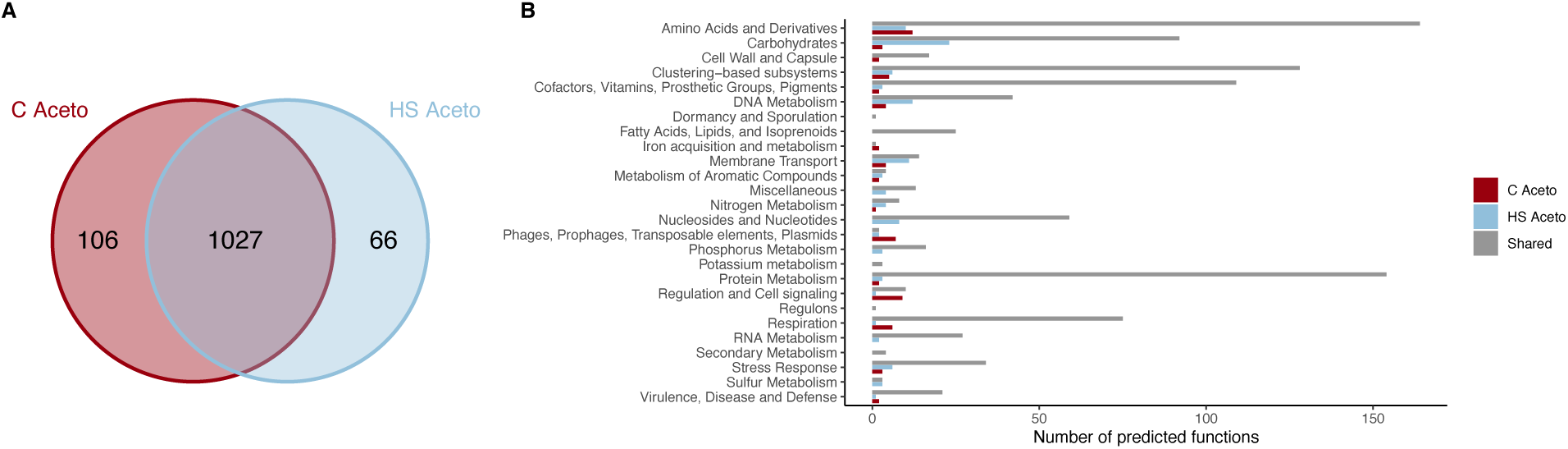
Summary of the RAST annotation for the C and HS Acetobacter. A) Venn diagram showing the number of functions unique to C Aceto (106/1133 total), unique to HS Aceto (66/1093 total), or shared (1027). B) Number of predicted functions per category colored by either unique to C Acetobacter (red), unique to HS Acetobacter (blue), or shared (grey). In general, both genomes encoded similar functions, but small differences are observed across the different categories.

**Supp. Fig. 1:**
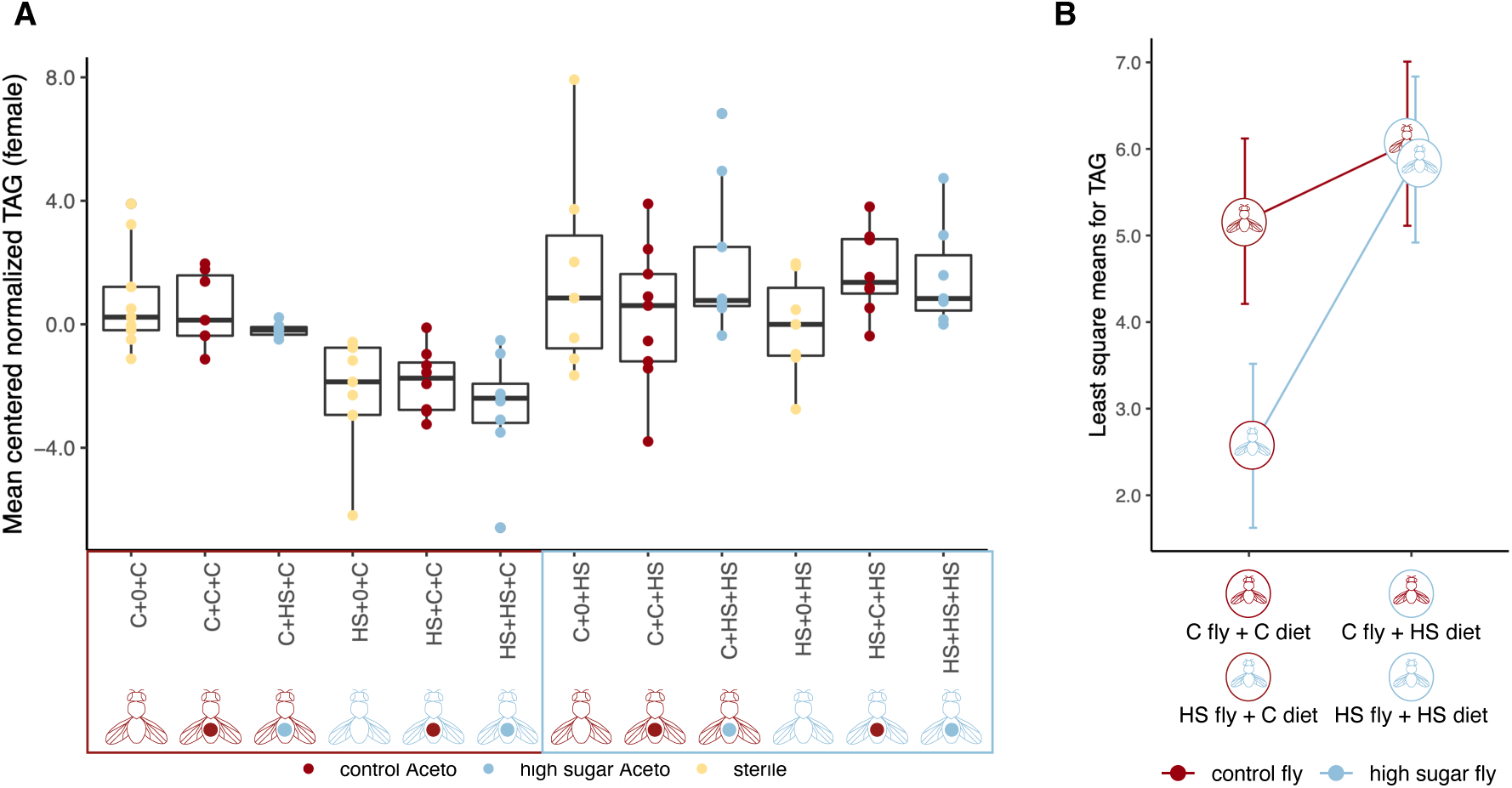
Metabolic effects of the *Acetobacter* transplant. Triglyceride levels were assessed from pools of three, age-matched (7-10 days old) females. Tryglycerides were measured using colorimetric assay test (Sigma TR0100-1KT) and then normalized to protein using a Bradford assay (BioRad 5000205). A) Boxplot for triglycerides (TAG) normalized to protein. TAG amount reflects fat storage in *Drosophila*. Each point represents a pool of three age-matched female flies. TAG levels were mean centered to control for each batch assayed. Interactions between fly genotype and diet shaped TAG levels (F_1,78.1_ = 9.00, p=0.004), but *Acetobacter* had no effect. B) LSM from the mixed model shows the significant interaction between fly genotype and diet. Error bars show standard error for LSM. TAG levels responded to diet for HS flies, but not for C flies.

## SUPPLEMENTARY TABLES

**Supp. Table 1:** Sample size and replication for development measured during the *Acetobacter* transplant experiment.

| Fly genotype | Microbiome | Diet | No. tubes | Avg larvae/tube | SE avg larvae/tube |
| --- | --- | --- | --- | --- | --- |
| control | sterile | control | 8 | 79.2 | 7.88 |
| control | sterile | high sugar | 5 | 55.2 | 11.1 |
| control | control | control | 4 | 92.2 | 10.1 |
| control | control | high sugar | 8 | 87.2 | 8.37 |
| control | high sugar | control | 5 | 85.2 | 5.89 |
| control | high sugar | high sugar | 11 | 78.3 | 4.6 |
| high sugar | sterile | control | 9 | 95.7 | 7.01 |
| high sugar | sterile | high sugar | 5 | 72.8 | 10.8 |
| high sugar | control | control | 11 | 84.7 | 9.44 |
| high sugar | control | high sugar | 6 | 108 | 9.21 |
| high sugar | high sugar | control | 12 | 87.3 | 10.9 |
| high sugar | high sugar | high sugar | 6 | 87 | 6.93 |

**Supp. Table 2:** Sample sizes for fecundity in the *Acetobacter* transplant experiment.

| Fly genotype | Microbiome | Diet | N Females |
| --- | --- | --- | --- |
| control | sterile | control | 44 |
| control | sterile | high sugar | 46 |
| control | control | control | 48 |
| control | control | high sugar | 71 |
| control | high sugar | control | 59 |
| control | high sugar | high sugar | 81 |
| high sugar | sterile | control | 45 |
| high sugar | sterile | high sugar | 45 |
| high sugar | control | control | 61 |
| high sugar | control | high sugar | 46 |
| high sugar | high sugar | control | 59 |
| high sugar | high sugar | high sugar | 59 |

**Supp. Table 3:** ANCOM results for differentially abundant ASVs from C and HS fly microbiomes. Taxonomy was characterized to the lowest level an assignment could be made, and most ASVs were classified to species level. CLR=centered log ratio, W=test statistic, Sig=significant difference between groups (N.S. is nonsignificant).

| ASV ID | CLR | W | Sig | Taxonomy |
| --- | --- | --- | --- | --- |
| d84f3e1a7dbb97bc4695b954c8a2d2b2 | 4.30129864 | 25 | HS fly | *Acetobacteraceae* |
| fb59adcbdd838dc7853f7388afb62b5a | 7.82830283 | 25 | HS fly | *Acetobacter pasteurianus* |
| 9329896cd08f88a88296d921369cd8ea | -3.4609509 | 20 | C fly | *Algoriella xinjiangensis* |
| f0d11f9f8ba452d5485a79e0b31b538f | -3.6539534 | 20 | C fly | *Acetobacter indonesiensis* |
| 3f2be3d57ac38f60b0b9c56fa5857f63 | -2.7057494 | 18 | N.S. | *Algoriella xinjiangensis* |
| 70e118808cdc81b5b2670837c7833d8d | -3.3688051 | 18 | N.S. | *Acetobacter persici* |
| dd4826376c1de49b8687c4f882d3bcf3 | -2.1106948 | 17 | N.S. | *Enterobacteriaceae* |
| 606aa8c7117f801dc5ccc8c2a78c68c3 | -1.7000214 | 16 | N.S. | *Commensalibacter* |
| 1c735b8a08dcdb5ca158987733234114 | -1.6719837 | 15 | N.S. | *Algoriella xinjiangensis* |
| da74ac9885f0aa2d29ec4338b46aa051 | 2.00310762 | 13 | N.S. | *Orbaceae* |
| 4043baca9447b9860efabab56603efc0 | 0.79635173 | 10 | N.S. | *Acetobacter species* |
| 61fd19eda2dee9c93613432491556a99 | 0.69392446 | 10 | N.S. | *Alphaproteobacteria* |
| 7070daee4850cf801719eab99ce88a82 | 0.96974134 | 10 | N.S. | *Commensalibacter* |
| b3ccbf1d4fc85411f948429591f371cf | 0.59968784 | 10 | N.S. | *Pseudomonas* |
| ca7279bfdcb977287193c567f96b025c | 0.75112887 | 10 | N.S. | *Acetobacter indonesiensis* |
| 0a04cded1e993516d5cdc851bfe07180 | -1.6048832 | 9 | N.S. | *Algoriella xinjiangensis* |
| 6e4961fe480153e97ebdcf8fca4913fb | 0.99515931 | 9 | N.S. | *Providencia* |
| 8dc187c9810636b7199c030caaaf0694 | 0.58697562 | 9 | N.S. | *Morganella* |
| 9f9bb5e1a63c5a0820d1571839534a2f | 0.62165362 | 9 | N.S. | *Commensalibacter* |
| 677d95c5a0bc1c25435861fe262cb487 | 0.50699766 | 8 | N.S. | *Alphaproteobacteria* |
| 4af26492bbe6d0d4980bc82c2f08f32d | 0.3940931 | 7 | N.S. | *Providencia* |
| ad8b60b9428bc76f3b50e428f4f5a29f | 0.41862229 | 7 | N.S. | *Stenotrophomonas* |
| f72576c61dde7b9b6ad683dea71b105f | 0.06411428 | 7 | N.S. | *Enterobacteriaceae* |
| 3c3c65e801e07d1be83bfa6d962b5627 | -0.3759816 | 6 | N.S. | *Commensalibacter* |
| 681de033774ada405895dd81abc7593f | -0.5750105 | 6 | N.S. | *Providencia rettgeri* |
| 97b76e52dafa224603888fb56ef0b831 | -0.3031253 | 5 | N.S. | *Commensalibacter* |

**Supp. Table 4:** Comparison of microbial community composition (Bray-Curtis dissimilarity) for recipients on C diets using PERMANOVA with 999 permutations.

| Category | Df | SS | R<sup>2</sup> | F | *P* value |
| --- | --- | --- | --- | --- | --- |
| Fly genotype | 1 | 0.683 | 0.021 | 5.220 | 0.007 |
| Microbiome | 1 | 12.415 | 0.380 | 94.938 | 0.001 |
| Fly genotype x Microbiome | 1 | 0.506 | 0.015 | 3.871 | 0.018 |
| Residual | 146 | 19.092 | 0.584 |  |  |
| Total | 149 | 32.695 |  |  |  |

**Supp. Table 5:**
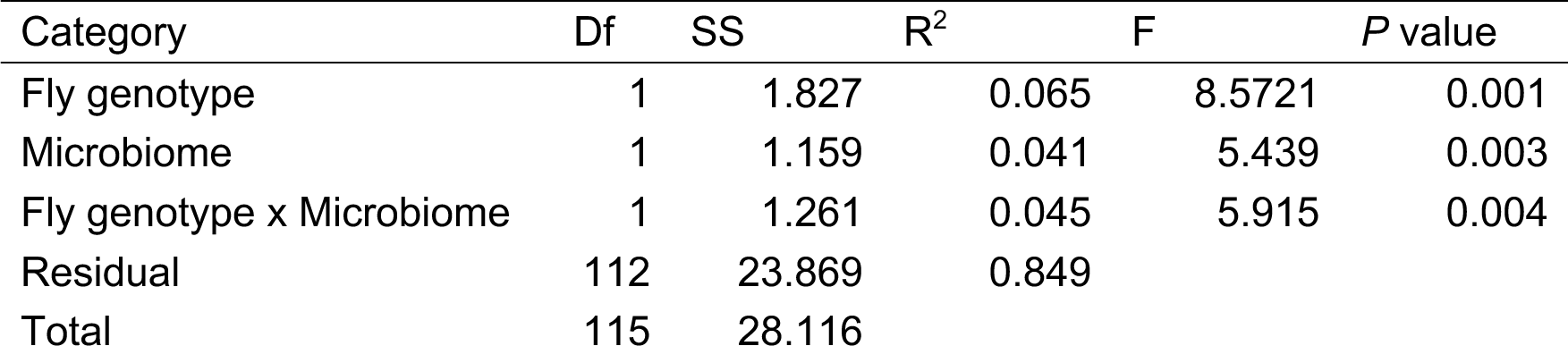
Comparison of microbial community composition for recipients on HS diets using PERMANOVA with 999 permutations.

**Supp. Table 6A:** Analysis of deviance for Cox model with interactions between fly genotype, microbiome, and diet for developmental timing in the *Acetobacter* transplant.

| Category | Log-likelihood | X<sup>2</sup> | Df | P value |
| --- | --- | --- | --- | --- |
| Null | -60640 |  |  |  |
| Fly genotype | -60584 | 111.642 | 1 | <2.20E-16*** |
| Acetobacter | -60517 | 134.874 | 2 | <2.20E-16*** |
| Diet | -58732 | 3568.585 | 1 | <2.20E-16*** |
| Fly x Acetobacter | -58675 | 115.244 | 2 | <2.20E-16*** |
| Fly x Diet | -58658 | 34.288 | 1 | 4.75E-09*** |
| Acetobacter x Diet | -58523 | 268.613 | 2 | <2.20E-16*** |
| Fly x Acetobacter x Diet | -58500 | 47.625 | 2 | 4.55E-11*** |

Significance codes: '\*\*\*' <0.001; '\*\*' <0.01; '\*' <0.05

**Supp. Table 6B:** Model selection statistics for developmental timing in the *Acetobacter* transplant. Model 1 is without interactions, while Model 2 includes all two and three-way interactions.

| Model | Log-likelihood | X<sup>2</sup> | X<sup>2</sup> Df | P(>\|Chi\|) |
| --- | --- | --- | --- | --- |
| Model 1 | -58732 |  |  |  |
| Model 2 | -58500 | 465.77 | 7 | <2.20E-16*** |

Significance codes: '\*\*\*' <0.001; '\*\*' <0.01; '\*' <0.05

**Supp. Table 7A:** Fixed effects for fecundity in the *Acetobacter* transplant. Significance was evaluated using Type III Wald F tests with Kenward-Roger degrees of freedom.

| Category | X<sup>2</sup> | Df | P value |
| --- | --- | --- | --- |
| Intercept | 637.8446 | 1 | <2.20E-16*** |
| Fly genotype | 4.0333 | 1 | 0.0446106* |
| Acetobacter | 39.7839 | 2 | 2.30E-09*** |
| Diet | 53.9716 | 1 | 2.03E-13*** |
| Fly x Acetobacter | 7.787 | 2 | 0.0203736* |
| Fly x Diet | 8.4003 | 1 | 0.0037517** |
| Acetobacter x Diet | 14.4492 | 2 | 0.0007285*** |
| Fly x Acetobacter x Diet | 16.3387 | 2 | 0.0002832*** |

Significance codes: '\*\*\*' <0.001; '\*\*' <0.01; '\*' <0.05

**Supp. Table 7B:**
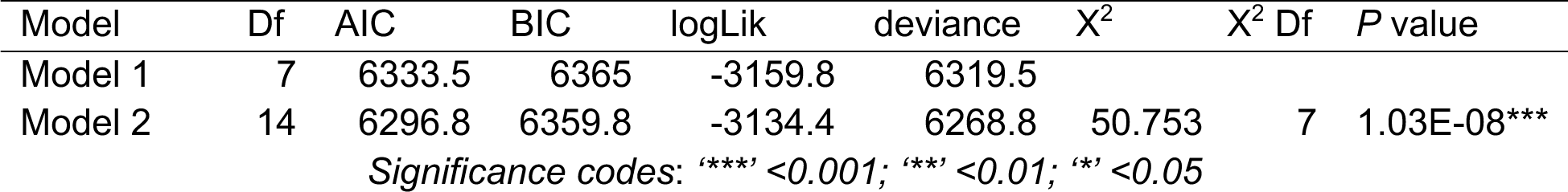
Model selection statistics for fecundity in the *Acetobacter* transplant. Model 1 is without interactions, while Model 2 includes all two and three-way interactions.

| Model | Df | AIC | BIC | logLik | deviance | X<sup>2</sup> | X<sup>2</sup> Df | P value |
| --- | --- | --- | --- | --- | --- | --- | --- | --- |
| Model 1 | 7 | 6333.5 | 6365 | -3159.8 | 6319.5 |  |  |  |
| Model 2 | 14 | 6296.8 | 6359.8 | -3134.4 | 6268.8 | 50.753 | 7 | 1.03E-08*** |

Significance codes: '\*\*\*' <0.001; '\*\*' <0.01; '\*' <0.05

**Supp. Table 8:** Average nucleotide identity (ANI) between C and HS *Acetobacter* strains.

| Metric | Value |
| --- | --- |
| ANI value (%) | 73.68 |
| C *Acetobacter* genome length (bp) | 3,436,380 |
| HS *Acetobacter* genome length (bp) | 2,895,396 |
| Average aligned length (bp) | 1,055,729 |
| C *Acetobacter* coverage (%) | 30.72 |
| HS *Acetobacter* coverage (%) | 37.10 |

**Supp. Table 9:** Uric acid degradation presence/absence in the most abundant bacteria from the C and HS microbiomes. Annotation was performed using RAST, and each were compared to the HS *Acetobacter* (Y= present, N= not present). Only HS *Acetobacter* encoded for uric acid degradation pathways.

| Role | HS Aceto | C Aceto | A. persici | Algoriella |
| --- | --- | --- | --- | --- |
| Urea ABC transporter, ATPase protein UrtD | Y | N | N | N |
| Urea ABC transporter, ATPase protein UrtE | Y | N | N | N |
| Urea ABC transporter, permease protein UrtB | Y | N | N | N |
| Urea ABC transporter, permease protein UrtC | Y | N | N | N |
| Urease accessory protein UreE | Y | N | N | N |
| Urease accessory protein UreF | Y | N | N | N |
| Urease accessory protein UreG | Y | N | N | N |
| Urease alpha subunit (EC 3.5.1.5) | Y | N | N | N |
| Urease beta subunit (EC 3.5.1.5) | Y | N | N | N |
| Urease gamma subunit (EC 3.5.1.5) | Y | N | N | N |

**Supp. Table 10:**
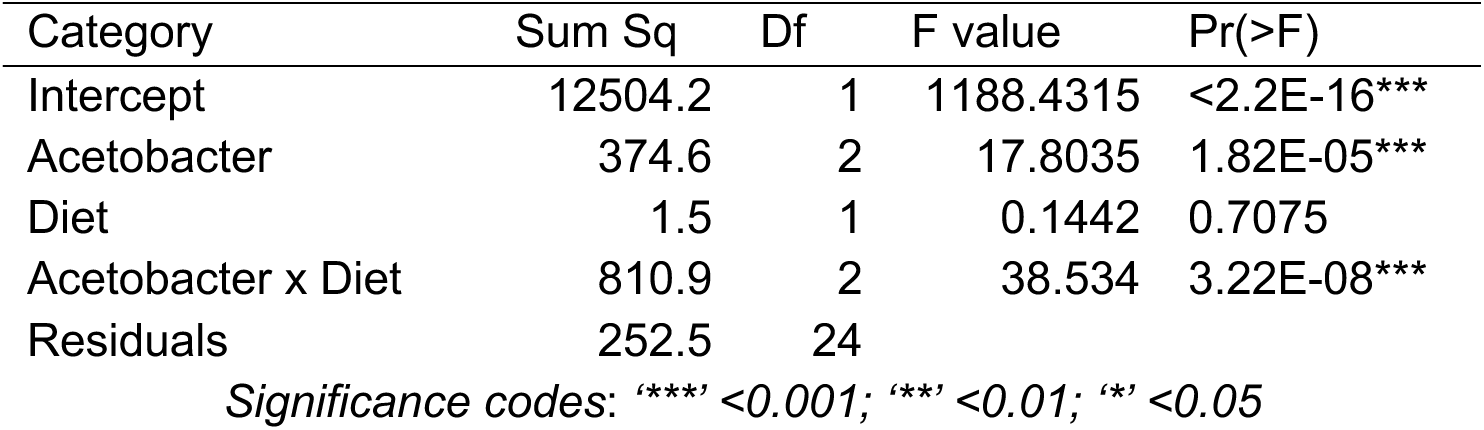
ANOVA results for uric acid degradation by *Acetobacters*.

| Category | Sum Sq | Df | F value | Pr(>F) |
| --- | --- | --- | --- | --- |
| Intercept | 12504.2 | 1 | 1188.4315 | <2.2E-16*** |
| Acetobacter | 374.6 | 2 | 17.8035 | 1.82E-05*** |
| Diet | 1.5 | 1 | 0.1442 | 0.7075 |
| Acetobacter x Diet | 810.9 | 2 | 38.534 | 3.22E-08*** |
| Residuals | 252.5 | 24 |  |  |

Significance codes: '\*\*\*' <0.001; '\*\*' <0.01; '\*' <0.05

